# Bone-Inspired Microarchitectured Materials with Enhanced Fatigue Life

**DOI:** 10.1101/494633

**Authors:** Ashley M. Torres, Adwait A. Trikanad, Cameron A. Aubin, Floor M. Lambers, Marysol Luna, Clare M. Rimnac, Pablo Zavattieri, Christopher J. Hernandez

## Abstract

Microarchitectured materials achieve superior mechanical properties through geometry rather than composition ^1-4^. Although lightweight, high-porosity microarchitectured materials can have high stiffness and strength, stress concentrations within the microstructure can cause flaw intolerance under cyclic loading ^5,6^, limiting fatigue life. However, it is not known how microarchitecture contributes to fatigue life. Naturally occurring materials can display exceptional mechanical performance and are useful models for the design of microarchitectured materials ^7,8^. Cancellous bone is a naturally occurring microarchitectured material that often survives decades of habitual cyclic loading without failure. Here we show that resistance to fatigue failure in cancellous bone is sensitive to the proportion of material oriented transverse to applied loads – a 30% increase in density caused by thickening transversely oriented struts increases fatigue life by 10-100 times. This finding is surprising in that transversely oriented struts have negligible effects on axial stiffness, strength and energy absorption. The effects of transversely oriented material on fatigue life are also present in synthetic lattice microstructures. In both cancellous bone and synthetic microarchitectures, the fatigue life can be predicted using the applied cyclic stress after adjustment for apparent stiffness and the proportion of the microstructure oriented transverse to applied loading. In the design of microarchitectured materials, stiffness, strength and energy absorption is often enhanced by aligning the microstructure in a preferred direction. Our findings show that introduction of such anisotropy, by reducing the amount of material oriented transverse to loading, comes at the cost of reduced fatigue life. Fatigue failure of durable devices and components generates substantial economic costs associated with repair and replacement. As advancements in additive manufacturing expand the use of microarchitectured materials to reusable devices including aerospace applications, it is increasingly necessary to balance the need for fatigue life with those of strength and density.

---

Fatigue failure is caused by the accumulation of microscopic damage following repeated loading and generates substantial economic costs associated with repair and replacement of durable devices. Microarchitectured materials are particularly susceptible to fatigue failure because the complex geometry can result in local stresses an order of magnitude greater than stresses applied to the bulk material ^5,6,9^. The presence of large stress concentrations can promote damage initiation and accumulation under cyclic loading and thereby reduce the fatigue life. Naturally occurring microarchitectured materials also experience cyclic loading and provide a model for strategies to resist fatigue failure.

Bone is a biological material with high stiffness and strength relative to density. In humans, most bones survive more than 50 years of habitual loading without failure. Whole bones consist of an outer shell made of dense tissue known as cortical bone that surrounds a foam-like tissue known as cancellous bone. Cancellous bone consists of a network of interconnected plate-like and rod-like struts called trabeculae (∼50-300 μm in thickness). Trabeculae in cancellous bone are preferentially aligned in the direction of stresses generated by habitual physical activity, resulting in a transversely isotropic microstructure. The fatigue life of cancellous bone and other cellular solids follows a normalized stress v. life (S-N) relationship under cyclic compressive loading ^10^:

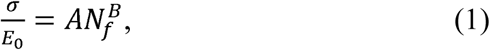

where σ is the maximum compressive stress, E_0_ is the initial Young’s modulus, N_f_ is the number of cycles to failure, and A and B are empirical constants.

To better understand the effects of microarchitecture on fatigue life, we determined the relationship between microstructure and fatigue damage processes in high porosity (> 90%) cancellous bone from the vertebral bodies of deceased human donors (n=44 specimens from 18 donors, see Methods). Cyclic compressive loading (zero to compression) was applied in the direction of habitual loading in vivo. Fatigue loading was suspended at a specified amount of cyclic loading (determined by accumulated cyclic strain) and the resulting amount and location of damage within the microstructure was detected using contrast agents (Fig. 1a and 1b, see Methods)^11,12^. Microarchitecture was assessed using three-dimensional images and analyzed using a morphological decomposition approach that isolates each individual strut within the structure and classifies the strut as plate-like or rod-like as well as determining its orientation relative to loading (Fig. 1c, d, see Methods)^13^. The amount of tissue damage caused by fatigue loading was correlated with maximum applied apparent strain (Extended Data Table S1) but was not correlated with specimen density or other specimen-average measures of microstructure. Surprisingly, the amount of tissue damage was reduced in specimens with thicker rod-like struts (Fig. 1e, Extended Data Table S2, R^2^=0.76, p < 0.01). This finding was unexpected since rod-like struts in cancellous bone constitute, on average, only 20% of the solid volume of high porosity cancellous bone (Extended Data Table S3) are primarily oriented transversely, carry only a small proportion of longitudinally-oriented loads, and therefore have negligible effects on stiffness and strength in the longitudinal direction ^14^ (transversely oriented elements have similar, small effects in nanolattice structures ^15^).

**Fig. 1.**
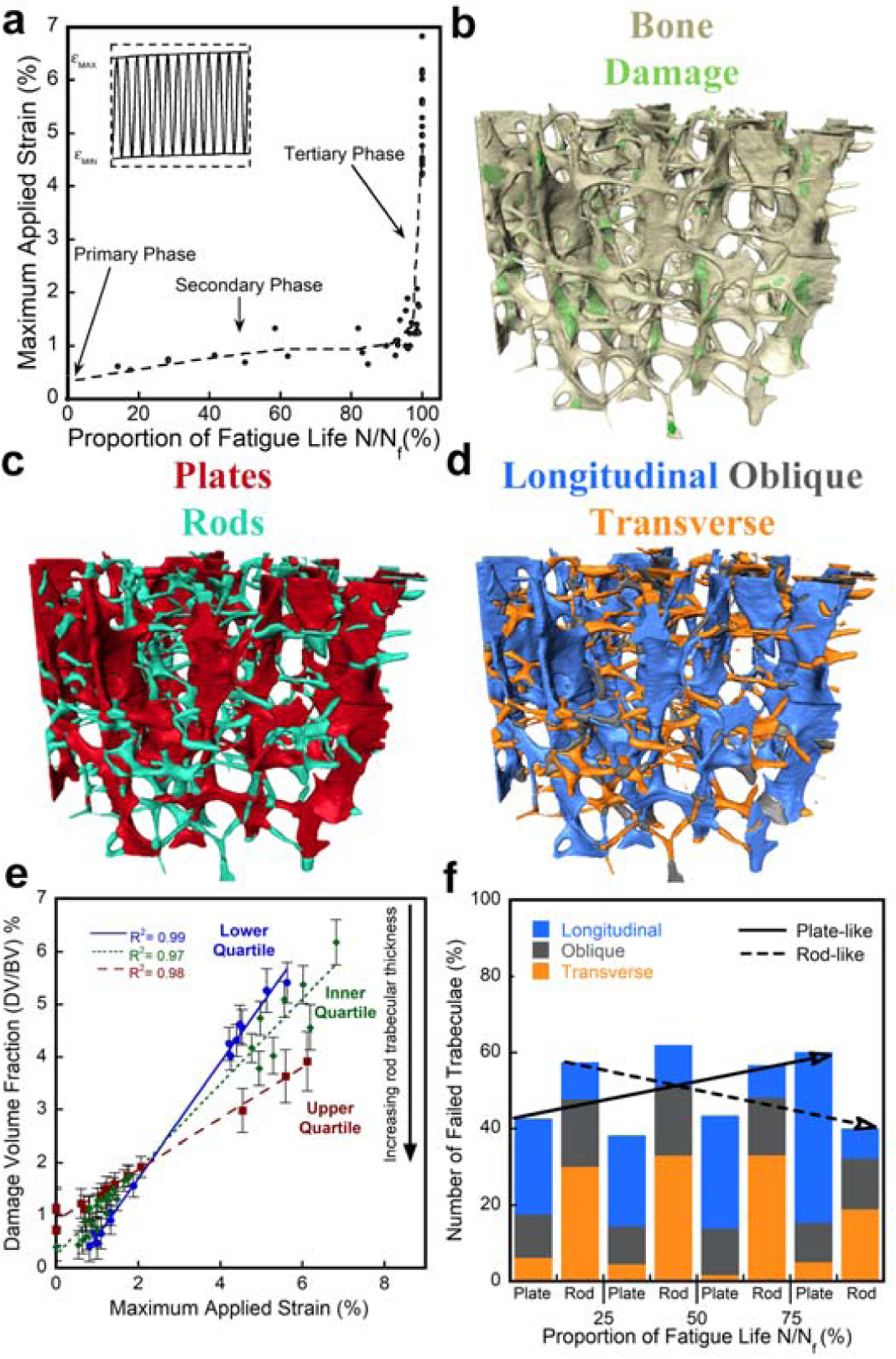
Microarchitecture influences fatigue damage accumulation in cancellous bone. **a,** The creep-fatigue curve of cancellous bone is shown with the three phases of fatigue loading indicated. Cyclic compressive loading of cancellous bone was stopped at different points along the creep fatigue curve (data points) to determine patterns of damage accumulation. Three dimensional images of cancellous bone with **b,** green indicating damage, **c,** plate-like and rod-like struts and **d,** strut orientation relative to anatomical position (longitudinal, oblique, and transverse). **e,** The amount of damage in cancellous bone was correlated with maximum applied strain, but specimens with thicker rod-like trabeculae experienced less damage accumulation (R^2^=0.76, p < 0.01). Error bars indicate the standard deviation as determined from the linear mixed effects model. **f.** Early in fatigue life, strut failure occurs primarily in transversely oriented rod-like struts; final mechanical failure is characterized by widespread failure of longitudinally oriented plate-like struts.

To better understand the effect of rod-like struts on fatigue failure we examined the distribution of tissue damage at different points during the fatigue loading process. Early in fatigue, failure of struts occurs primarily in rod-like trabeculae; substantial damage accumulation in plate-like trabeculae does not occur until overt failure (Fig. 1f). The pattern of strut failure is also related to orientation, failed rod-like trabeculae are predominately transversely oriented while failed plate-like trabeculae are predominately oriented longitudinally (Extended Data Fig. S1). The experimental findings were consistent with finite element models that indicated that apparent compressive loading results in tensile stresses in rod-like trabeculae (primarily transversely oriented) and compressive stresses in plate-like trabeculae (primarily longitudinally oriented, Extended Data Fig. S2). These findings suggest that, in cancellous bone, transversely oriented trabeculae act as sacrificial elements during cyclic loading by accumulating tissue damage and thereby protecting the load carrying, longitudinally oriented, plate-like trabeculae.

Tissue heterogeneity is also a major contributor to damage accumulation in cancellous bone ^12,16^ and is therefore a potential explanation for our findings in human bone tissue. To confirm that the effects of transversely oriented struts was due to geometry rather than material heterogeneity we generated three dimensional models of cancellous bone microstructures using a high-resolution projection stereolithography printer (Fig. 2 a,b, M1, Carbon, USA) and a urethane methacrylate resin (UMA) ^17^. Cancellous bone microstructures (Fig. 2b, n=5) were modified by adding material to the surface of transverse trabeculae in one of three increments: no modification (original geometry), +20 μm on the surface (an average increase in rod thickness of 20% ± 5%, mean ± SD), or +60 μm on each surface (an average increase in rod thickness of 45% ± 14%). Because transverse rod-like trabeculae constitute only a small portion of the solid volume and carry only a small portion of loads due to their orientation, thickening of rod-like struts had only a small effect on density (Fig. 2c, increase of 11% ± 8%, mean ± SD) and apparent stiffness (22% ± 19% increase in longitudinal Young’s modulus, Fig. 2d). However, these small increases in thickness of rod-like trabeculae led to increases in fatigue life by as much as two orders of magnitude (Fig. 2e, Extended Data Fig. S3).

**Fig. 2.**
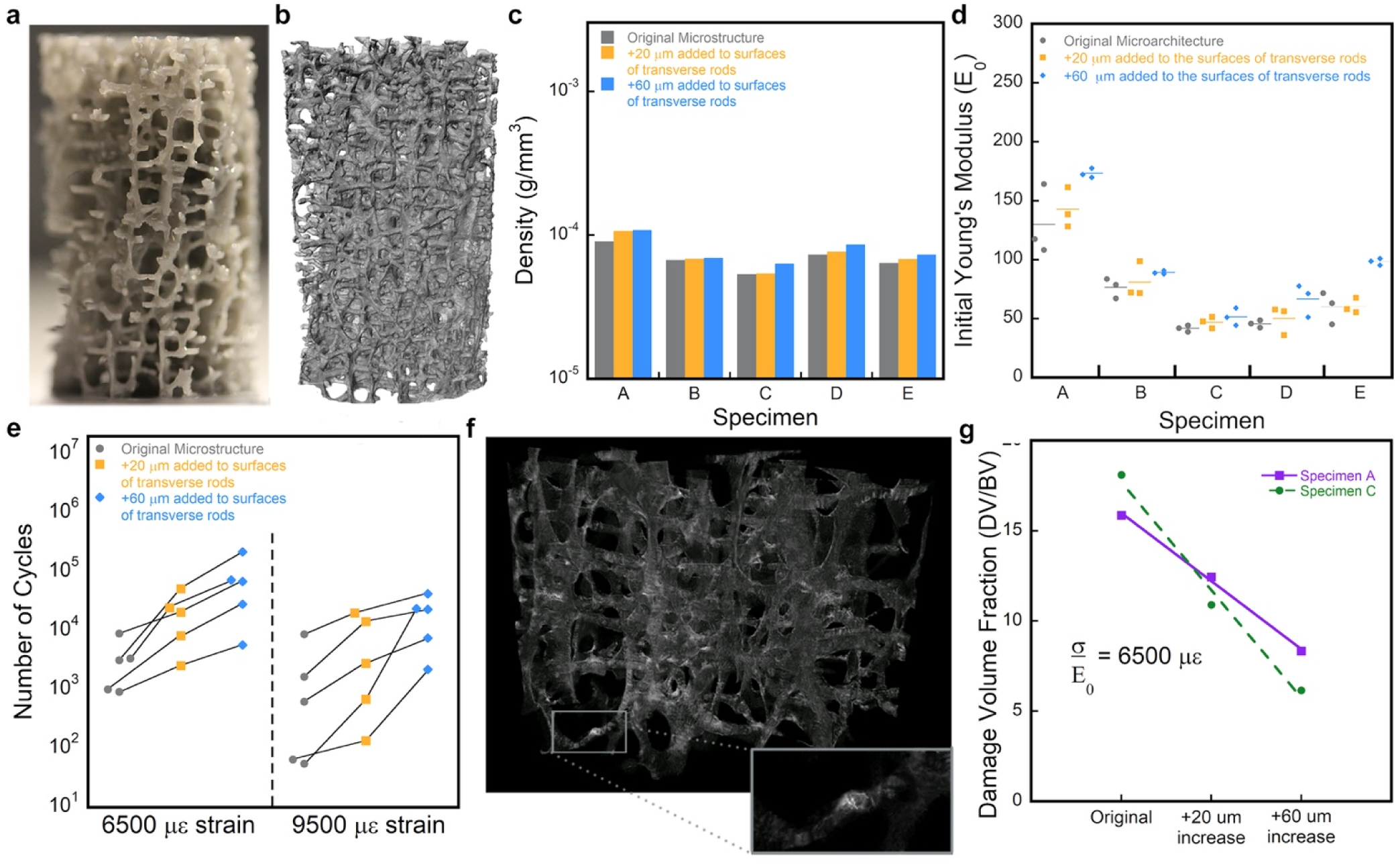
| Models of cancellous bone generated using additive manufacturing show that fatigue life is sensitive to small changes in microarchitecture. **a,** digital images of human vertebral cancellous bone were edited and printed into **b.** high-resolution three-dimensional models. Increases in the thickness rod-like struts had small effects on **c,** density, and **d.** stiffness (Young’s modulus in first cycle of loading), yet resulted in **e,** increases in fatigue life by as much as 2 orders of magnitude (lines connect samples derived from the same bone specimen, two magnitudes of normalized cyclic stress are shown). **f,** A micro-computed tomography image of a three-dimensional printed sample of cancellous bone after fatigue loading to failure. A radio-opaque dye penetrant indicated regions of accumulated damage. **g,** The amount of damage generated by fatigue loading to failure was reduced in 3D printed specimens with greater thickness of rod-like struts.

To confirm that damage accumulation in the 3D printed models was similar to that of cancellous bone, we examined damage in the microstructure using a radio-opaque dye penetrant (Fig. 2f). Printed specimens with thicker rod-like struts showed reduced damage accumulation (Fig. 2g). Hence, localization of failure in the printed models matched the patterns seen in cancellous bone, suggesting similar failure mechanisms among materials and further supporting the idea that increases in fatigue life are due to changes in microarchitecture and not material heterogeneity. Furthermore, finite element models of the specimens indicated that the average stresses in rod-like trabeculae (predominately transversely oriented) were greater than those in plate-like trabeculae (predominately longitudinally oriented, Extended Data Fig. S2), demonstrating the localization of damage follows load distributions within the microarchitecture. Together these findings indicate that small increases in mass applied to transversely oriented structural components of the microstructure can reduce matrix stresses leading to disproportionately large, beneficial effects on fatigue life.

To determine if the effects of transverse elements on fatigue failure are generalizable to other cellular solids, we created printed models of an octet truss ^1^ as well as an octet truss modified to have plate-like and rod-like elements mimicking the microstructure and anisotropy of cancellous bone (Fig. 3a). The octet truss shows a stretching dominated deformation behavior, cancellous bone microstructure shows bending dominated behavior and the bone-like microarchitecture displayed a combination of both stretching and bending deformation behaviors (Extended Data Fig S4). In the bone-like microarchitectures, increases in transverse strut thickness resulted in an increase in the fatigue life by a factor of eight (Fig. 3b) with only a small change in density (+4%) or longitudinal stiffness (+20%, Extended Data Table S4). In the octet truss, increases in transverse strut thickness resulted in an increase in fatigue life by a factor of five (Fig. 3b) with only minor changes in density (+10%) or uniaxial stiffness (+14%, Fig. 3b, Extended Data Table S4). In contrast, when the orientation of the octet trusses with thickened elements was rotated by 90 degrees so that thickened elements were oblique to the applied loads, the fatigue life was *reduced* by a factor of 9 (Fig. 3b), demonstrating that the effect of transverse struts on fatigue life is related to the proportion of material oriented transverse to loading rather than the thickness of the transverse struts *per se*. To understand the extent to which the transversely oriented material influenced fatigue damage accumulation we performed nonlinear finite element models of cyclic loading. Fatigue damage involves a local irreversible energy dissipating process resulting in increases in inelastic dissipation energy. Finite element models of cyclic loading indicate that the fatigue life of the octet and bone-like microarchitectures with and without thickened struts is closely related to the inelastic dissipation energy per unit work (Fig. 3c). Hence, increases in the transverse volume fraction (Ψ, the proportion of the solid volume oriented transverse to loading) in these microarchitectured materials reduce the amount of inelastic energy dissipation and damage accumulation during cyclic loading, just as thicker rod-like trabeculae (predominately transversely oriented) experienced less damage accumulation in cancellous bone (Fig 1e). Together these findings show that the effects of transverse struts on fatigue life is not unique to cancellous bone but extends to synthetic microarchitectured materials. That our findings regarding damage accumulation are consistent in human bone tissue (a biological ceramic polymer composite) as well as a polymer used in additive manufacturing further suggests that the effect is due to geometry and is not limited to one class of constituent material.

**Fig. 3.**
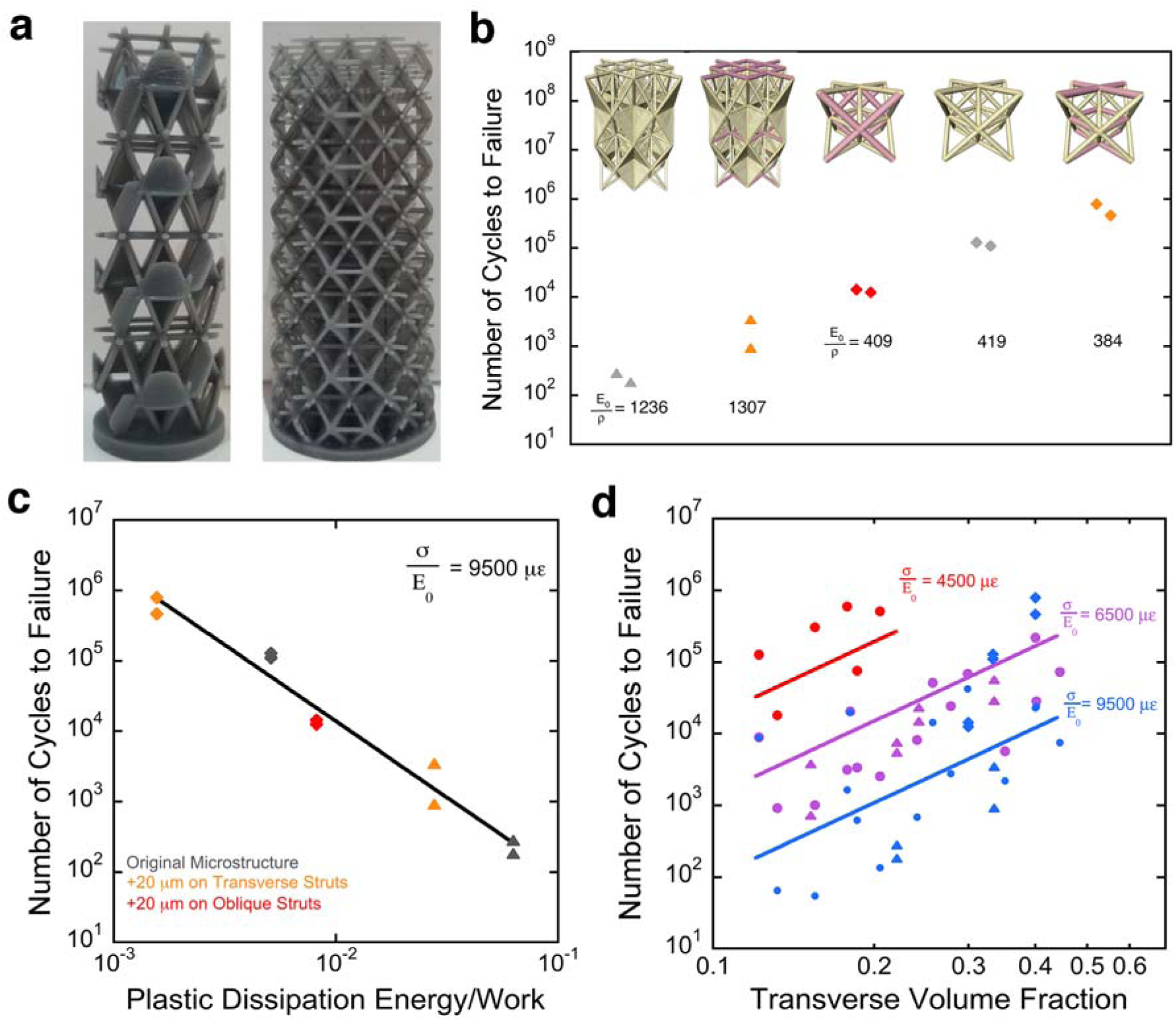
Transverse volume influences fatigue life in repeating cellular solids. **a,** Images of the bone-inspired microstructure and an octet truss are shown. **b,** The fatigue life of microarchitectured materials printed as designed or with rod-like struts thickened (colored) is shown. Thickening transverse struts increases fatigue life, while thickening vertically oriented struts reduces fatigue life (specific stiffness, E_0_/ρ is also shown). **c,** Fatigue life of the lattice structures is related to the inelastic dissipation energy per unit work determined from finite element models. **d,** Fatigue life for 3D printed specimens of bone (•), bone-like microstructure (▴) and octets (♦) at different applied cyclic loading (σ/E_0_, noted in με). Lines indicate the regression model fits (Equation 2, R^2^ = 0.82).

We developed empirical models to characterize the relationship between fatigue life (N_f_), applied cyclic normalized stress (noted as σ/E_0_ for cellular solids), and transverse volume fraction (Ψ) for bone, bone-like microarchitectures and octet trusses. Surprisingly, the regression models identified a predictive equation only slightly different from (1):

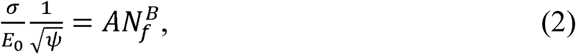

in which A and B are empirical constants (R^2^ = 0.82, Fig. 3d, Extended Data Table S5). This simple modification to the normalized S-N relationship provides a means of considering fatigue life during the design of microarchitectured materials. Designs with greater proportions of struts aligned axial to loading are more efficient in terms of specific stiffness and strength ^15^, but will also experience reduced fatigue life due to reductions in the proportion of material oriented transverse to expected loading. Future applications of high porosity microarchitectured materials to durable products such as vehicles will require balancing performance in terms of stiffness, strength and energy absorption with costs associated with replacement and repair due to fatigue failure.

**Supplementary Information** is linked to the online version of the paper

## Acknowledgments

The authors thank Rob Shepherd for comments.

## Funding

This work was supported in part by the National Institute of Arthritis and Musculoskeletal and Skin Diseases of the National Institutes of Health (U.S) under Award Numbers AR057362 and AR073454; the National Science Foundation (NSF) through a Graduate Research Diversity Supplement (GRDS) to NSF 1068260 and an NSF GRFP (AMT), and CAREER award CMMI 1254864; as well as a Cornell-Colman Fellowship (AMT). Imaging data was performed in the Cornell BRC-Imaging Facility (NIH S10OD012287). The content of the work is solely the responsibility of the authors and does not necessarily represent the official views of the funding agencies.

## Author contributions

AMT, FML, CMR, PZ and CJH designed the study. AMT, FML, ML and CAA performed experiments and image acquisition and analysis. AAT and PZ performed computational analyses. AMT and CJH drafted the manuscript. All authors provided critical review of the raw data and the final manuscript.

## Competing interests

The authors declare no competing interests.

Correspondence and requests for materials should be addressed to

## Methods

### Mechanical Characterization and Damage Assessment in Cancellous Bone

This study examined human vertebral cancellous bone (n=44 cancellous bone specimens from 10 male and 8 female donors, 62-92 years of age years of age) as reported in two prior studies ^11,12^. Cyclic loading was applied to a specified increase in cyclic strain and expressed as the average portion of the fatigue life (N/N_f_) using a previously identified empirical model (Fig. 1a).

Samples of human vertebral cancellous bone were acquired from the lumbar vertebrae of deceased donors (tissue source NDRI). The donors had no history of metabolic bone disease or cancer. The biomechanical analysis included cancellous bone from the third lumbar vertebral bodies of 16 donors ^11^ and from the fourth lumbar vertebral bodies of 12 donors ^12^. Cylindrical specimens, 8 mm in diameter and 27 mm in length were collected in the cranial-caudal direction using a coring tool. Two specimens were collected from each L3 vertebra and one specimen was collected from each L4 vertebrae resulting in a total of 44 specimens. Specimens were wrapped in saline soaked gauze and stored in airtight tubes at −20 °C until mechanical testing. Bone marrow was removed with a low-pressure water jet.

Mechanical characterization of high porosity cellular solids can be challenging as the individual struts of the foam deform locally at the contact with loading platens resulting in end artifacts that generate non-uniform strains within the gage length ^18^. To avoid end artifacts, specimens were press fit into cylindrical brass end-caps and secured with cyanoacrylate glue (Loctite 401, Newington, CT, USA). Using this approach, loads are transferred to the specimen via shear at the specimen/endcap interface thereby avoiding end artifacts. After being secured in end caps the bone specimens were stored overnight at 4 °C and hydrated with saline soaked gauze to allow the glue to fully cure.

Specimens were placed in a servo hydraulic testing machine (858 Mini Bionix, MTS, Eden Prairie, MN, USA) and loaded in cyclic compression at room temperature (23 °C) to induce damage. During testing, hydration was maintained by placing a rubber membrane around the specimen containing physiologically buffered saline (pH of 7.4). Strain was measured with a 25 mm extensometer (MTS, Eden Prairie, MN, USA) attached to the specimen’s end-caps and force was measured with a load cell (100 lb capacity, SSM-100, Transducer Techniques, CA, USA). Prior to each bout of loading, ten preconditioning cycles between 0 and 0.1% strain at a rate of 0.5% per second was applied. Fatigue loading was applied using a 4 Hz haversine waveform cyclically between 0 N and a compressive load corresponding to σ = E_0_* 0.0035 mm/mm, where σ is stress and E_0_ is the initial Young’s modulus of the specimen (determined from the preconditioning cycles). Force and strain data were collected to calculate Young’s modulus (E), reduction in Young’s modulus (1 - E/E_0_), and maximum strain ^11,12^.

Specimens from the third lumbar vertebrae included five specimens that were not subjected to fatigue loading ^11^. Fatigue loading of the remaining specimens was stopped before failure by monitoring the creep-fatigue curve, secant modulus, and hysteresis loops to stop loading at a predetermined magnitude of cyclic strain. Fatigue loading was stopped at the start of the secondary phase, in the middle of the secondary phase, and at different points within the tertiary phase. Six specimens from the third lumbar vertebrae were loaded well beyond the start of the tertiary phase to 4% apparent strain. Specimens were then stained for damage using lead-uranyl acetate. The number of cycles to failure (N_f_) was determined for specimens loaded to an apparent strain of 4%. For the remaining specimens, the number of cycles to failure (N_f_) was estimated using an empirical relationship between reduction in Young’s modulus and proportion of fatigue life ^11^.

Specimens from the fourth lumbar vertebrae were loaded in two separate bouts of cyclic loading at room temperature (23 °C) ^12^. The first bout of fatigue was stopped at the beginning of the tertiary phase (Fig.1a), identified as a the rapid increase in accumulation of apparent strain ^11^. Following the first bout of cyclic loading, specimens with end caps were carefully removed and bulk stained in xylenol orange solution (0.5 mM, Sigma Chemical Co., St. Louis, MO) to stain damage caused by the first bout of loading. The specimens were carefully returned to the material testing device, and a second bout of cyclic loading was applied, until the specimens reached 5% apparent strain ^11^. Specimens were carefully removed from the testing device and bulk stained in calcein solution (0.5 mM, Sigma Chemical Co., St. Louis, MO) to stain damage generated during the second bout of loading. Specimens were then embedded undecalcified in methyl methacrylate.

Damage accumulation in specimens from the third lumbar vertebral bodies was determined using the lead uranyl acetate stain and x-ray microcomputed tomography (10 μm isotropic voxels). A global threshold was used to identify regions of bone tissue and damage stain (see ^11^). Damage generated in specimens from the fourth lumbar vertebrae was determined using images obtained through serial milling (for a complete review of serial milling procedure, we direct the reader to prior work ^19,20^). Serial milling achieved a 0.7 × 0.7 × 5.0 μm voxel size image of bone and each of the fluorescent markers of microdamage. Tissue damage stained and visualized using these approaches includes both microscopic cracks as well as regions of constrained microcracking (known as diffuse damage in the bone literature). As described in prior studies, the amount of tissue damage was correlated with the maximum applied strain during cyclic loading. The relationship between maximum applied strain and amounts of microdamage was similar for both staining/visualization techniques (Extended Data Fig. S5).

### Cancellous Bone Microarchitecture

Microarchitecture was assessed using specimen average measurements (BoneJ, bonej.org, ^21^, 21 μm isotropic voxel microcomputed tomography images collected prior to fatigue loading, Supplementary Information Table S6) as well as morphological analysis of each individual trabecula (Supplementary Information Table S7). Morphological analysis involved classifying each trabecula as rod-like or plate-like based on Digital Topological Analysis (DTA) (Fig. 1C, ITS software, Columbia University). Additionally, the orientation of individual trabeculae was determined as the angle from the superior-inferior direction and classified as longitudinal (0 ≤ β ≤ 30°), oblique (30 < β ≤ 60°), and transverse (60 < β ≤ 90°) (Fig.1d) ^13,22^.

The amount of damage on plate- or rod-like trabeculae and the amount of damage present on trabeculae with each orientation (longitudinal, oblique or transverse) was determined using custom scripts written for use with Matlab (Mathworks, Natick, MA, USA) and Amira (5.3 Visage Imaging, San Diego, CA, USA).

Failure of cancellous bone has been referred to as accumulation of failed trabeculae ^23,24^. Here we consider failure of a trabecula to occur when at least 10% of the volume of the trabecula includes microdamage ^25^. To understand patterns in failure of individual trabeculae, we report the proportion of failed plate-like trabeculae (# failed plate-like trabeculae/ # plate-like trabeculae), and the proportion of failed rods (Extended Data Fig. S1, #failed rod-like trabeculae/ # rod-like trabeculae).

### Additive Manufacturing of Modified Cancellous Bone Microstructures

Microcomputed tomography images of a subset of cancellous bone samples (n=5, 10 μm voxel images, 8 mm diameter, ∼20mm length) ^11,12^ that had been collected before mechanical loading were modified digitally by adding material to the surface of transverse rod-like trabeculae to three different amounts: +60 μm (+120 μm strut thickness, 45% ± 14% increase in rod thickness), +20 μm (+40 μm in rod thickness, 20% ± 5% increase in strut thickness) or no modification (original geometry). A high-resolution stereolithography system (M1, Carbon, USA) was used to generate three-dimensional models of the modified and unmodified microstructures from a urethane methacrylate polymer resin (UMA 90, Carbon, USA, E = 2 GPa, ^17^) at 1.5 times isotropic magnification (12 mm diameter, ∼30 mm length). The accuracy of the printed geometries was confirmed through direct comparison of the three-dimensional .stl image to a microcomputed tomography image of the printed model (Supplementary Information Table S7. Morphological analysis of each individual trabecula was analyzed for the three-dimensional printed specimens (Supplementary Information Table S8).

Mechanical characterization of models of cancellous bone generated through additive manufacturing was performed as described above. Specimens were submitted to cyclic fatigue loading from zero to a normalized initial compressive stress magnitude σ/E_0_ corresponding to 9500 με, 6500 με or 4500 με until failure (4% applied strain, Extended Data Fig. S3).

Damage in polymer specimens was identified using a radio opaque dye penetrant. Samples were soaked for 24 hours at room temperature (23 °C) in the dye penetrant containing 250 g zinc iodide, 80 ml distilled water, 80ml isopropyl alcohol and 1 ml Kodak photo solution ^26^. Samples were left to dry for 12 hours to remove any excess dye prior to image acquisition using x-ray microcomputed tomography (10 μm) ^27^.

### Design and Manufacturing of Bone-like Architectured Material

Bone-like microarchitectures derived from the octet truss were designed in ABAQUS/CAE. The length of the octet truss was elongated along one axis (the longitudinal axis) by 40% so that struts were oriented 60 degrees from the transverse plane, thereby mimicking longitudinal elements seen in cancellous bone. To further match the architecture of the cancellous bone, 6 plates were added per unit octet. Transversely isotropic supercells were created from unit cells in which plates were included in either the upper or lower half of each unit cell (Fig. S4). The resulting supercell achieved porosity, transverse volume fraction, longitudinal volume fraction and plate volume fraction similar to the cancellous bone specimens. These “supercell” structures were then exported as .stl files from the ABAQUS/CAE sketcher and used for the 3D printing process.

### Finite Element Modeling

Simulations were performed using ABAQUS/Standard. Models were meshed with C3D8 brick elements (1.5 million/model) and quasi-static analyses were carried out for cyclic compressive loading consistent with the same strain that was applied during experiments. The material was assumed to be elastic-perfectly plastic with an initial Young’s Modulus of 600 MPa and a yield stress of 26 MPa. The structure was loaded under displacement-control along the loading axis with rollers on the other end and free boundary conditions on all other sides. Simulations were run for 5 to 25 compression loading-unloading cycles. Simulations were performed on a high performance computing cluster consisting of 10-core Intel Xeon-E5 processors. Inelastic dissipation was extracted from each model at the end of all loading cycles.

To characterize the primary deformation mechanism within each model (bending dominated or stretching dominated) the stress triaxility was determined at each point within the model. Stress triaxility is defined as:

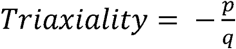

where *p* is the hydrostatic stress, defined as positive when stress is compressive and *q* is the vonMises stress. By this definition, points under uniaxial tension would have a triaxiality of +0.33, and points under pure uniaxial compression would have a triaxiality value of −0.33. Histograms summarizing the values of stress triaxiality within microstructures (Fig S4) under longitudinal compression show the deformation behavior for each architecture. As expected, the octet shows only stretching deformation (large peaks only at +0.33 or −0.33). The cancellous bone microstructure shows a high amount of bending behavior (uniform distribution of triaxiality throughout with peaks at +0.33 and −0.33) and the bone-like architectures show a combination of bending and stretching deformation behavior.

### Statistical Treatment

A multivariate correlation analysis was used to identify trends between measures of damage, changes in mechanical properties and volume-averaged measures of bone microarchitecture (Extended Table S1). Others have shown that the amounts of microdamage formed during fatigue loading was associated with maximum applied strain and reduction in Young’s modulus and was not related to volume-averaged measures of microarchitecture ^11^. A backward elimination was performed using a linear mixed model with fixed effects to identify aspects of trabecular microstructure that were associated with the accumulation of fatigue damage. Donor was included as a random effect to take into account the use of multiple specimens from each donor.

To examine the pattern of failure of discrete trabeculae observed during fatigue loading, an ANCOVA was used to test for differences in the relationship between reductions in Young’s modulus and the number of failed plate- and rod-like trabeculae. An ANOVA was used to test for differences in the number of failed trabeculae in each orientation (longitudinal, oblique, transverse) and post-hoc comparisons were performed using Tukey HSD test.

A Generalized Least squares Model (GLM) was used to identify the empirical relationship between stress amplitude (σ/E_0_), the transverse volume fraction (Ψ), and number of cycles to failure (N_f_, Equation 2). To account for differences in the number of samples for each microstructure (1-2 specimens per microstructure) microstructure was included as a random effect using REML. Statistical tests were conducted using JMP 12 (SAS Institute Inc., Cary, NC, USA).

## Data and materials availability

The data that support the findings of this study are available from the corresponding author upon reasonable request.

