## Supplementary material for "Bone-Inspired Microarchitectured Materials with Enhanced Fatigue Life"

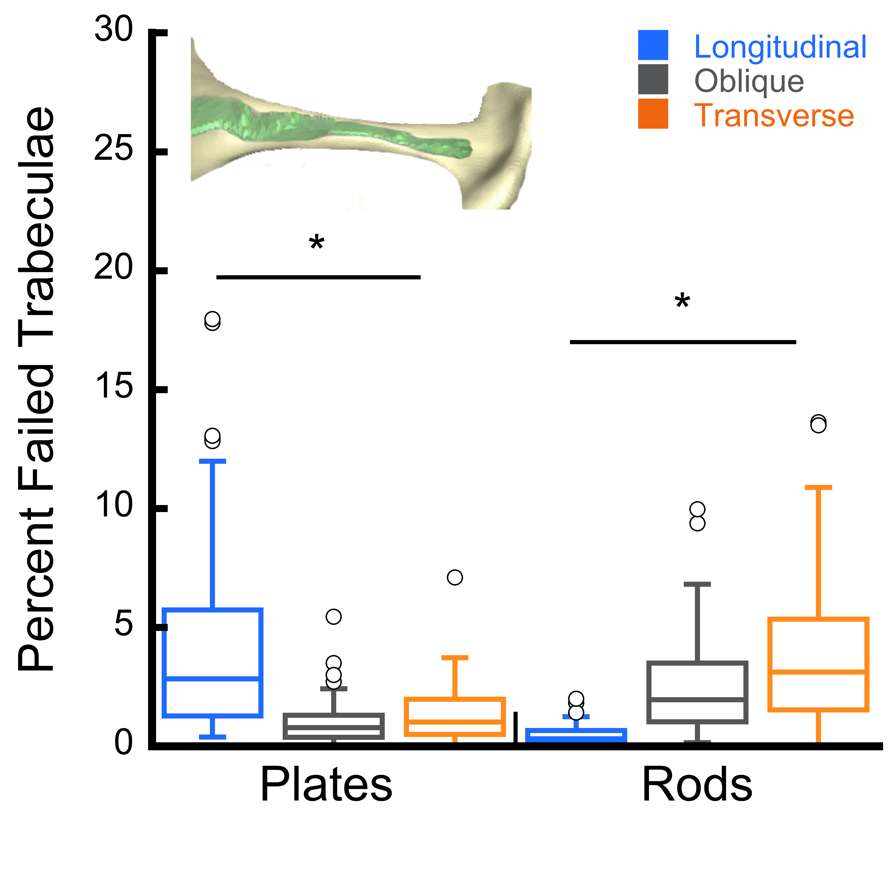

**Extended Data** **Fig. S1 The proportion of failed plate and rod-like trabeculae are shown by orientation.** Failure of individual trabeculae occurred primarily in longitudinally oriented plate-like struts and transversely oriented rod-like struts (*, p < 0.01). Inset: A failed rod-like strut (damage is indicated by green color).

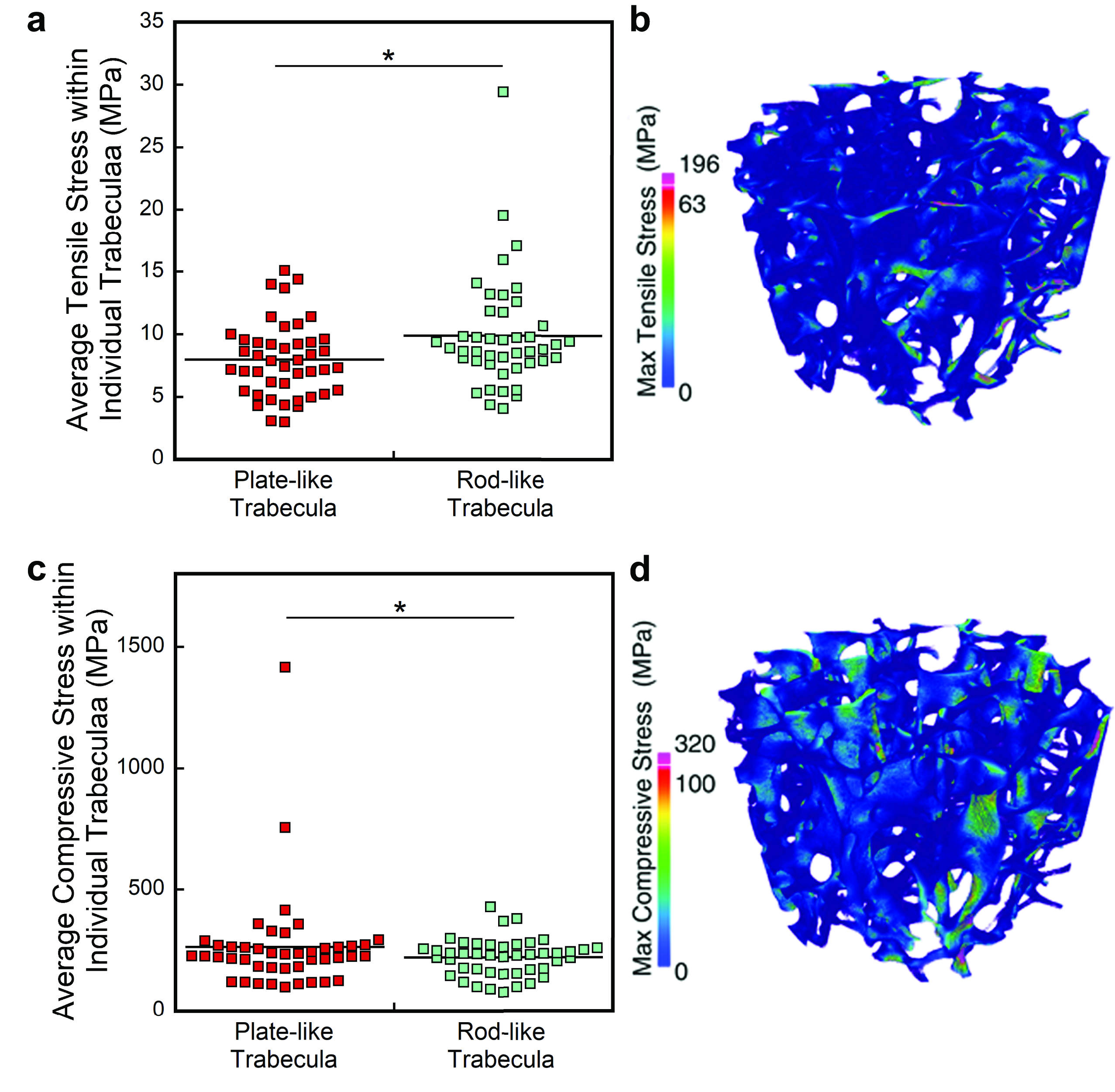

**Extended Data** **Fig. S2. The average stress per strut is shown. a,b** Tensile stresses are greater in rod-like (primarily transversely oriented) trabeculae. **c,d** Compressive stresses are greater in plate-like (predominately longitudinally oriented) trabeculae. *, p < 0.01.

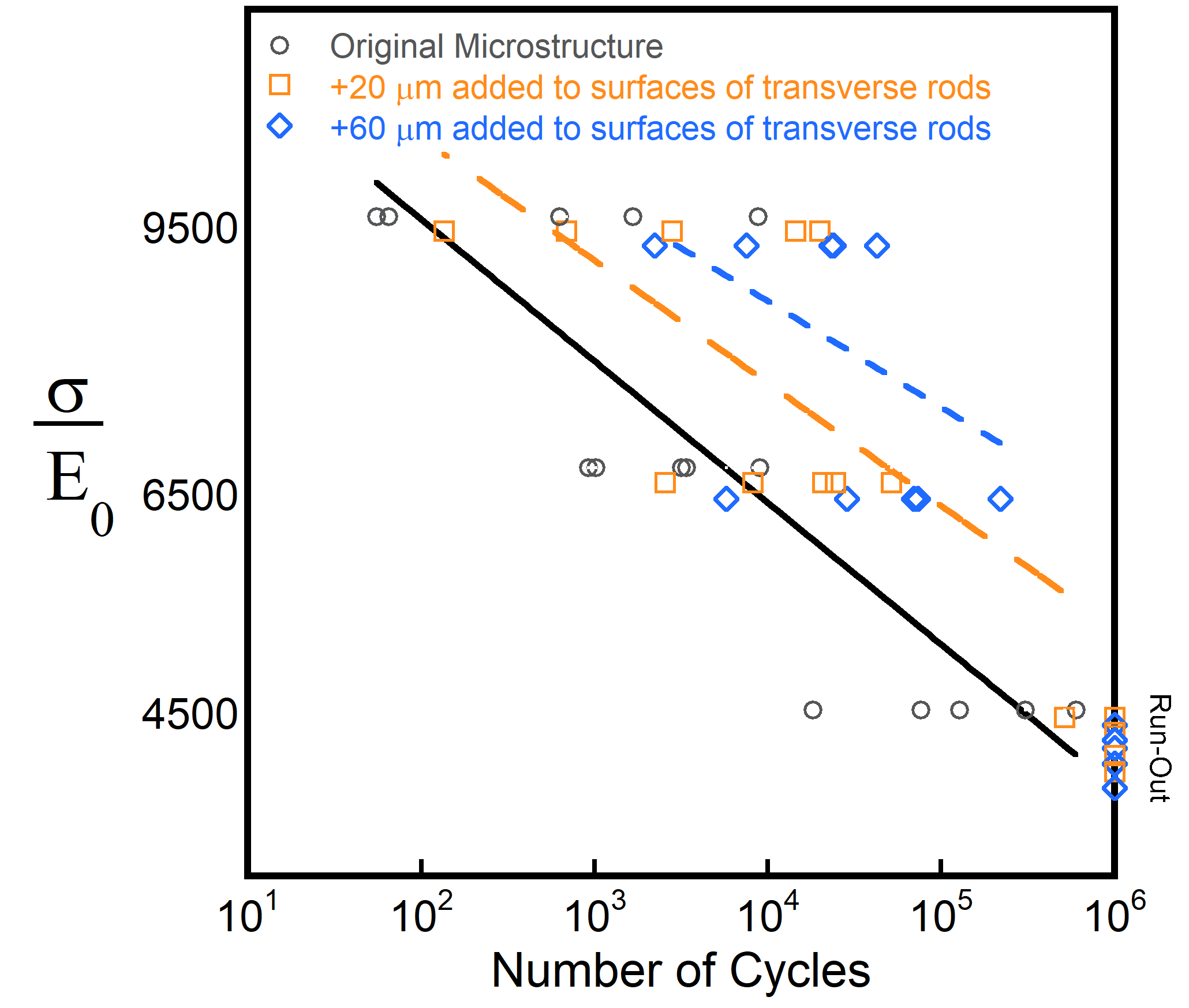

**Extended Data** **Fig. S3.** The number of cycles to failure increased in specimens with increased rod-thickness (on average 10-100 times). Gray represents specimens with the original microarchitecture, orange and blue represent specimens with a 20 μm and 60 μm added thickness to the surface of the transverse rod-like trabeculae. Run-out was defined at 10^6^ cycles.

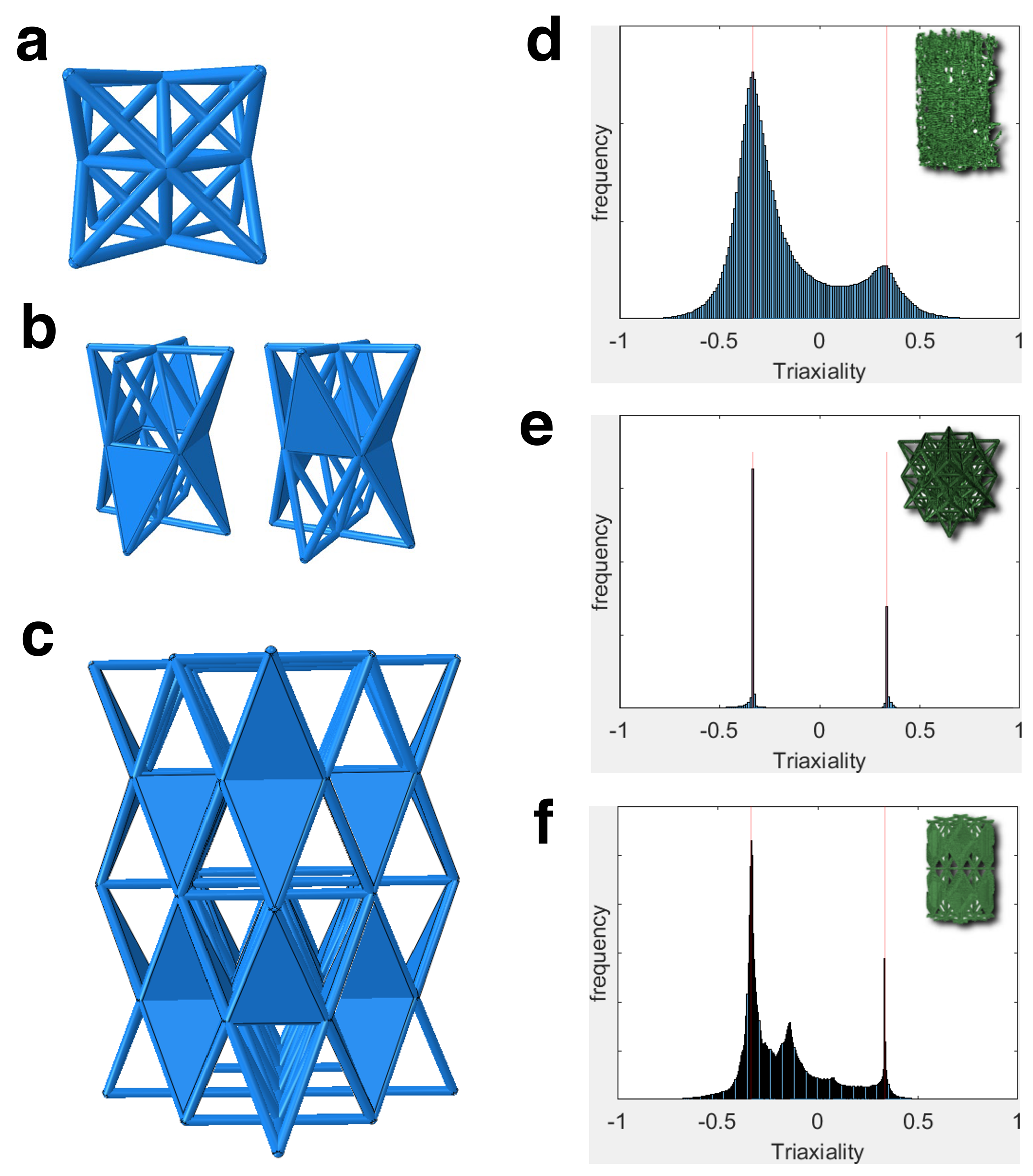

**Extended Data** **Fig. S4 | a.** the octet truss. **b.** two bone-like unit cells that are combined into **c.** a 2 X 2 X 2 supercell bone-like architecture. Stress triaxiality for each microstructure (inset are the meshes for each structure) with vertical red lines indicating +0.33 and -0.33 values for **d.** Cancellous bone, **e**. Octet and **f.** Bone-like microstructure.

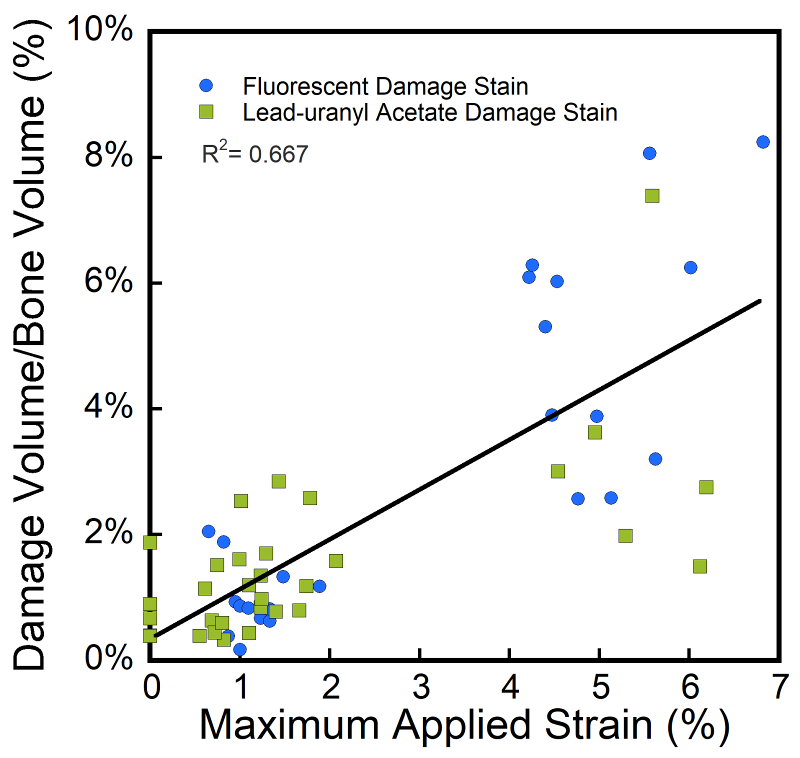

**Extended Data** **Fig. S5 | The two staining methods achieved similar relationships between amount of tissue damage and applied apparent strain.** Validation of the two methods of staining microscopic damage. Blue points represent fluorescently stained samples that were imaged three-dimensionally with serial-milling, and green points represent lead-uranyl acetate stained samples imaged with x-ray microcomputed tomography.

**Table S1. Damage accumulation was positively correlated with maximum applied strain.** The lower triangle displays the correlation coefficients and the 95% confidence interval is shown in the upper triangle. *p < 0.05

|  | Reduction in  Young's  modulus (%) | Maximum  Applied  Strain (%) | Damage  Volume  Fraction (%) | Bone  Volume  Fraction  (%) | Trabecular  Thickness  (µm) | Degree of  Anisotropy | Structural  Model  Index |
| --- | --- | --- | --- | --- | --- | --- | --- |
| Reduction  in  Young's  modulus (%) | - | (0.77, 0.92) | (0.42, 0.75) | (-0.43, 0.08) | (-0.14, 0.38) | (-0.20, -0.32) | (-0.01, 0.48) |
| Maximum  Applied  Strain (%) | 0.86* | - | (0.64, 0.86) | (-0.34, 0.18) | (-0.09, 0.42) | (-0.53, -0.05) | (-0.04, 0.46) |
| Damage  Volume  Fraction  (%) | 0.61* | 0.77* | - | (-0.33, 0.19) | (0.00, 0.49) | (-0.57, -0.10) | (-0.01, 0.49) |
| Bone  Volume  Fraction  (%) | -0.19 | -0.09 | -0.07 | - | (-0.13, 0.39) | (-0.14, 0.38) | (-0.72, -0.34) |
| Trabecular  Thickness  (µm) | 0.13 | 0.17 | 0.26 | 0.14 | - | (-0.63, -0.20) | (0.18, 0.62) |
| Degree of  Anisotropy | -0.06 | -0.31 | -0.36 | 0.13 | -0.44 | - | (-0.76, -0.42) |
| Structural  Model  Index | 0.25 | 0.22 | 0.26 | -0.56 | 0.42 | -0.62 | - |

**Table S2. Damage was correlated with rod trabecular thickness.** A linear mixed effects model for predicting damage volume fraction and the parameters are listed in the table (R^2^ =0.76, n=44).

| **Parameter** | **Coefficent** | **SE** | **DF Den** | **t Ratio** | **p** |
| --- | --- | --- | --- | --- | --- |
| Maximum Applied Strain (%) | 0.77 | 0.08 | 42 | 9.66 | <0.0001* |
| Rod Trabecular Thickness | -9.88 | 9.16 | 48 | -1.08 | 0.0685 |
| Maximum Applied Strain (%) * Rod Trabecular Thickness | -12.50 | 4.38 | 43 | -2.85 | 0.0066* |

**Table S3.**

Plate- and rod-like microarchitecture of cancellous bone (n=44).

| **Microarchitecture Measure** | **Mean ± SD** |
| --- | --- |
| Plate bone volume fraction (pBV/TV, %) | 7.53 ± 2.92 |
| Rod bone volume fraction (rBV/TV, %) | 2.06 ± 0.87 |
| Axial bone volume fraction (aBV/TV, %) | 6.36 ± 2.17 |
| Plate tissue fraction (pBV/BV, %) | 78.15 ± 6.44 |
| Rod tissue fraction (rBV/BV, %) | 21.85 ± 6.44 |
| Plate trabecular number (pTb.N, 1/mm) | 2.89 ± 0.21 |
| Rod trabecular number (rTb.N, 1/mm) | 2.35 ± 0.58 |
| Trabecular plate thickness (pTb.Th, mm) | 0.12 ± 0.02 |
| Trabecular rod diameter (rTb.Th, mm) | 0.12 ± 0.02 |
| Trabecular plate surface area (pTb.S, mm^2^) | 0.09 ± 0.02 |
| Trabecular rod length (rTb.*l*, mm) | 0.45 ± 0.03 |
| Rod-Rod Junction density (R-R Junc. D, 1/mm^3^) | 1.26 ± 0.48 |
| Plate-Rod Junction density (R-P Junc. D, 1/mm^3^) | 6.24 ± 2.21 |
| Plate-Plate Junction density (P-P Junc. D, 1/mm^3^) | 5.54 ± 2.30 |

**Table S4.** Microarchitectual parameters and mechanical properties of octet truss and bone-like repeating pattern.

| Unit Cell | Number  of  Rods | Number  of  Plates | Porosity  (%) | Transverse  Strut  Thickness  (mm) | Density  (g/mm^3^) | $\frac{\boldsymbol{\sigma}}{\boldsymbol{E}_{\boldsymbol{0}}}$ | Initial  Young's  Modulus  (MPa) | Number  of  Cycles to  Failure  (N) |
| --- | --- | --- | --- | --- | --- | --- | --- | --- |
| Octet Truss | 36 | 0 | 92.56 | 0.400 | 4.22E-05 | 9500 | 31.21  ±1.40 | 120309  ±12659 |
| Octet Truss  +20 μm  Transverse Rods | 36 | 0 | 90.71 | 0.534 | 4.66E-05 | 9500 | 35.69  ±1.82 | 628980  ±232017 |
| Octet Truss  +20 μm  Oblique Rods | 36 | 0 | 90.71 | 0.534 | 4.66E-05 | 9500 | 38.03  ±0.32 | 13532  ±1251 |
| Bone-like  Repeating  Pattern | 20 | 6 | 87.99 | 0.400 | 6.13E-05 | 9500 | 148.54  ±16.73 | 233  ±69 |
| Bone-like Repeating  Pattern +20 μm  Transverse Rods | 20 | 6 | 87.07 | 0.534 | 6.35E-05 | 9500 | 169.08  ±10.06 | 2210  ±1830 |
| Bone-like  Repeating  Pattern | 20 | 6 | 87.99 | 0.400 | 6.13E-05 | 6500 | 137.46  ±16.19 | 6550  ±1445 |
| Bone-like Repeating  Pattern +20 μm  Transverse Rods | 20 | 6 | 87.07 | 0.534 | 6.35E-05 | 6500 | 174.05  ±1.85 | 43045  ±19668 |

**Table S5.** **Regression models reveal an adjustment to the S-N relationship to account for the effects of transverse volume.** Regression equations are displayed in the form Log N_f_ = a_0_ + a_1_*Log(σ/E_0_) + a_2_*Log(ψ), R^2^ = 0.82, p < 0.001.

| **Coefficient** | **Value** | **S.E.** |
| --- | --- | --- |
| a_0_ | 76.0 | 6.0 |
| a_1_ | -6.9 | 0.6 |
| a_2_ | 3.5 | 0.5 |

**Supplementary Information**

Title: Bone-Inspired Microarchitectured Materials with Enhanced Fatigue Life

**Authors:** Ashley M. Torres^1,2^, Adwait A. Trikanad^3^, Cameron A. Aubin^1^, Floor M. Lambers^1^, Marysol Luna^1^, Clare M. Rimnac^4^, Pablo Zavattieri^3^, Christopher J. Hernandez^1,2,5*^

**Table S6.**

Traditional bone microarchitecture measurements of cancellous bone (n=44).

| **Microarchitecture Measure** | **Mean ± SD** |
| --- | --- |
| Bone volume fraction (BV/TV, %) | 8.13 ± 2.16 |
| Bone surface (BS, mm^2^) | 520.54 ± 226.68 |
| Bone surface to bone volume ratio (BS/BV, mm^2^/mm^3^) | 20.09 ± 3.09 |
| Trabecular thickness (Tb. Th, µm) | 131.74 ± 17.34 |
| Trabecular separation (Tb. Sp, µm) | 1240.50 ± 301.28 |
| Degree of anisotropy (DA) | 1.56 ± 0.18 |
| Structure model index (SMI) | 1.63 ± 0.36 |
| Connectivity density (Conn. D, mm^-3^) | 2.73 ± 1.25 |

**Table S7. Accuracy of three-dimensional printed geometries was confirmed with microCT images of each specimen.** Number and p

late- and rod-like trabeculae of three-dimensional specimens of cancellous bone (n=5).

| **Specimen** | **Increase in Transverse Rods** | **Number of Rods** | **Number of Plates** | **Bone Volume Fraction (BV/TV)** | **microCT measured BV/TV from 3D printed samples** | **Percent Error in Additive Manufacturing (AD, %)** |
| --- | --- | --- | --- | --- | --- | --- |
| A | Original | 4873 | 9414 | 0.1567 | 0.1451 | 7.42 |
| A | 20 micron | 4372 | 9552 | 0.1586 | 0.1480 | 6.69 |
| A | 60 micron | 4358 | 9645 | 0.1925 | 0.1766 | 8.25 |
| B | Original | 2743 | 5228 | 0.1126 | 0.1120 | 0.54 |
| B | 20 micron | 2252 | 2998 | 0.1223 | 0.1187 | 2.99 |
| B | 60 micron | 2185 | 3015 | 0.1411 | 0.1295 | 8.31 |
| C | Original | 2660 | 4897 | 0.0853 | 0.0780 | 8.67 |
| C | 20 micron | 2443 | 4889 | 0.0891 | 0.0836 | 6.17 |
| C | 60 micron | 2521 | 4924 | 0.1093 | 0.0996 | 8.96 |
| D | Original | 3922 | 6267 | 0.1016 | 0.0985 | 3.10 |
| D | 20 micron | 3674 | 6405 | 0.1063 | 0.1038 | 2.44 |
| D | 60 micron | 3877 | 6388 | 0.1359 | 0.1478 | 8.73 |
| E | Original | 3796 | 5514 | 0.0934 | 0.0849 | 9.12 |
| E | 20 micron | 3659 | 5600 | 0.1004 | 0.0936 | 6.85 |
| E | 60 micron | 3862 | 5589 | 0.1298 | 0.1179 | 9.13 |

**Table S8.** Plate- and rod-like microarchitecture of three-dimensional printed cancellous bone (n=5).

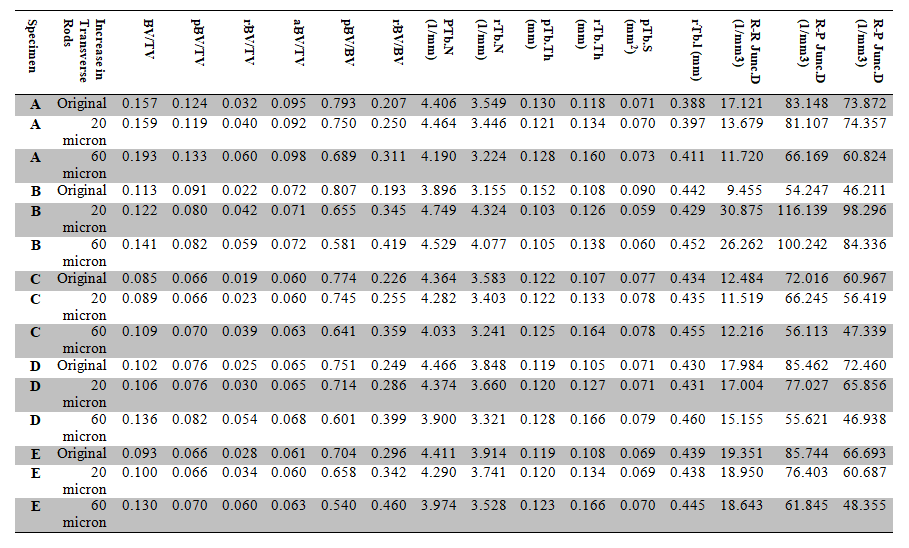
